# Structurally related but genetically unrelated antibody lineages converge on an immunodominant HIV-1 Env neutralizing determinant following trimer immunization

**DOI:** 10.1101/2021.04.09.439148

**Authors:** Safia S. Aljedani, Tyler J. Liban, Karen Tran, Ganesh Phad, Suruchi Singh, Viktoriya Dubrovskaya, Pradeepa Pushparaj, Paola Martinez-Murillo, Justas Rodarte, Alex Mileant, Vidya Mangala Prasad, Rachel Kinzelman, Sijy O’Dell, John R. Mascola, Kelly K. Lee, Gunilla B. Karlsson Hedestam, Richard T. Wyatt, Marie Pancera

## Abstract

Understanding the molecular mechanisms by which antibodies target and neutralize the HIV-1 envelope glycoprotein (Env) is critical in guiding immunogen design and vaccine development aimed at eliciting cross-reactive neutralizing antibodies (NAbs). Here, we analyzed monoclonal antibodies (mAbs) isolated from non-human primates (NHPs) immunized with variants of a native flexibly linked (NFL) HIV-1 Env stabilized trimer derived from the tier 2 clade C 16055 strain. The antibodies displayed neutralizing activity against the autologous virus with potencies ranging from 0.005 to 3.68 ug/ml (IC_50_). Structural characterization using negative-stain EM and X-ray crystallography identified the variable region 2 (V2) of the 16055 NFL trimer to be the common epitope for these antibodies. The crystal structures revealed that the V2 segment adopts a β-hairpin motif identical to that observed in the 16055 NFL crystal structure. These results depict how vaccine-induced antibodies derived from different clonal lineages penetrate through the glycan shield to recognize a hypervariable region within V2 (residues 184-186) that is unique to the 16055 strain. They also provide an explanation for the potent autologous neutralization of these antibodies, confirming the immunodominance of this site and revealing that multiple angles of approach are permissible for affinity/avidity that results in potent neutralizing capacity. The structural analysis reveals that the most negatively charged paratope correlated with the potency of the mAbs. The atomic level information is of interest to both define the means of autologous neutralization elicited by different tier 2-based immunogens and facilitate trimer redesign to better target more conserved regions of V2 to potentially elicit cross-neutralizing HIV-1 antibodies.

**Author summary:** NHPs immunizations with an HIV-1 immunogen (native-like tier 2 clade C 16055 strain) elicit potent HIV-1 tier 2 autologous polyclonal neutralizing antibodies. To understand the basis of the autologous neutralization, we determined structures of antibodies isolated from the vaccinated NHPs in complex with their epitopes. Our structural analysis reveals that the V2 hypervariable region, unique to 16055, is immunodominant and targeted by antibodies from diverse lineages. Additionally, vaccine-elicited V2 NAbs use different binding angles to avoid Env N-glycan shield and the more negatively charged paratope displays potent autologous neutralizing function. In summary, detailed analysis of how vaccine-elicited monoclonal antibodies interact with the target antigen provide valuable information for the design of immunogens aimed to elicit more broadly HIV-neutralizing antibodies. The use of cocktail/prime-boost sequential regimens that include a range of sequence variation combined with the removal/shielding of unwanted immunodominant epitopes will likely be needed to reach this goal.

## Introduction

The human immunodeficiency virus type 1 (HIV-1) is one of the major health challenges with 38 million cases worldwide [1, 2]. The numerous HIV-1 strains are classified into four groups: M, N, O, and P, based on their zoonotic transmission history [3, 4]. Group M is responsible for the majority of HIV-1 infections worldwide and is further divided into at least nine genetically distinct clades: A, B, C, D, F, G, H, J, K, and circulating recombinant form (CRFs) based on their geographic distribution [1, 5]. The highest infection rate is in Southern Africa, India, and Ethiopia (clade C), which total 46% of the HIV-1 infections worldwide. Therefore, an HIV-1 vaccine aimed to protect against transmission of clade C variants is a prioritized goal [1].

To provide protective immunity against the diverse array of HIV-1 strains circulating in the human population, broadly neutralizing antibodies (bNAbs) targeting the conserved regions of the variable HIV-1 envelope glycoprotein (Env) spike are needed. The HIV-1 Env, the main target of bNAbs, is a heterotrimeric glycoprotein located at the surface of the virus [6]. So far, a vaccine capable of eliciting such responses has proven challenging due to the numerous immune escape properties the functional HIV-1 Env spike has evolved, including high antigenic diversity, heavy N-linked glycosylation, conformational masking and quaternary packing that occludes efficient antibody access to cross-conserved determinants [7–13]. Nonetheless, an HIV-1 vaccine against diverse isolates and in particular clade C strains that cause most disease has been the focus of many studies [4, 14–16].

Several designs of stabilized soluble Env trimers that mimic the functional viral spike were generated for clinical evaluation and vaccine development once near-atomic level structure of Env was obtained [8, 17–23]. A cleavage-independent near-native soluble Env mimic with native flexibly linked (NFL) trimer was successfully engineered for the clade C strain 16055 and its structure was determined [18, 19, 24]. Recent studies reported that immunization of rhesus macaques with the stabilized 16055 NFL TD CC “I201/A433C” induced serum antibody responses capable of neutralizing the 16055 autologous tier 2 virus. Furthermore, mAbs that mediated this activity were isolated and shown to bind a highly variable epitope determinant in the Env V2 region, as determined by alanine scanning mutagenesis and differential adsorption [25, 26].

To further understand the immune response to the stabilized clade C 16055 immunogen following different immunization strategies in NHPs, we characterized mAbs from different immunization groups and determined their interactions with their epitope at low and high resolution, using nsEM and/or X-ray crystallography, respectively. Interestingly, all the NAbs recognized the same neutralizing hypervariable V2 loop but use different V genes, reflected in their differences in chemistry (electrostatic potential) and different angles of approach. This indicates that the polyclonal immune response favors these immunodominant epitopes, which are unique to the strain used for immunogen design. Our structural analysis suggests that careful analysis of both the sequence and structure of immunogens should be taken into account for next generation vaccine design: this immunodominant loop could be deleted or glycan-masked in priming immunizations to potentially shift neutralizing responses to more conserved determinants to more efficiently elicit cross-neutralizing antibodies.

## Results

### Immunization with native-like tier 2 Clade C NFL trimers elicit potent tier 2 autologous neutralizing antibodies from a polyclonal serum pool

We have shown previously that the well-ordered, stabilized NFL Env trimer [19] elicited HIV-1 autologous tier 2 neutralizing Abs in NHP [25, 26]. Here, we analyzed mAbs isolated from different immunization strategies with variants of the NFL Env-stabilized trimer in Chinese rhesus macaques (*Macaca mulatta*) to better understand the specificity of the elicited immune response. The animals were immunized at week 0, 4 and 12 (**Fig 1A**). Group A was immunized with 16055 NFL trimers conjugated to liposomes [25], group B was immunized with soluble 16055 NFL trimers with glycans at N276, N301, N360 and N463 deleted (degly 4 (Δ276, Δ301, Δ360, and Δ463)) and group C was immunized with soluble 16055 degly 4 trimers at week 0 and 4 and boosted with the 16055 NFL with glycans restored (wild type, WT) at week 12 (**Fig 1B**). Two weeks after the third immunization, samples were collected. Plasma neutralization assays indicated that animals in all 3 groups developed tier 2 16055 autologous titers (**S1 Fig**).

**Fig 1.**
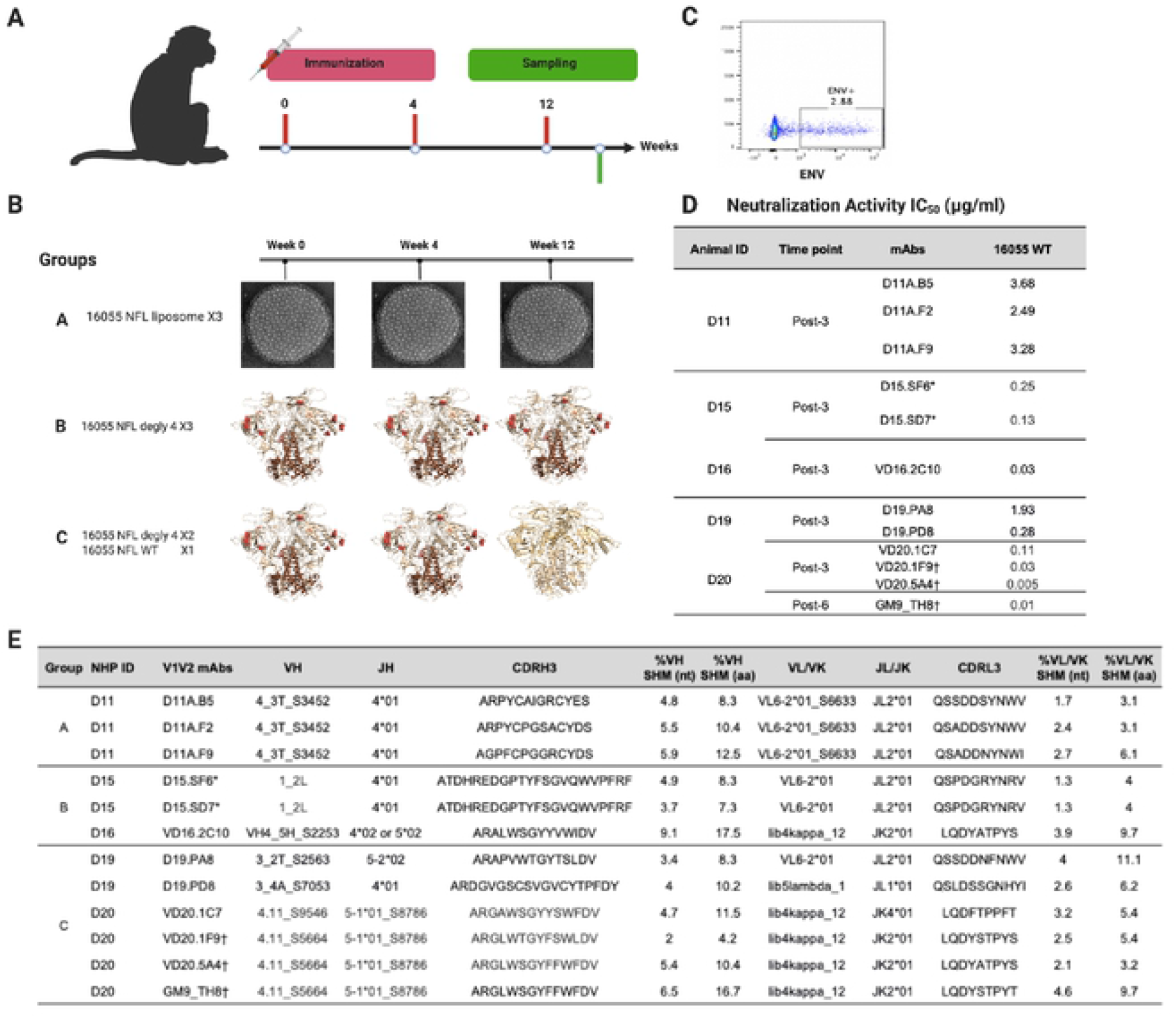
Immunization with 16055 Clade C NFL variants in NHP and autologous neutralization from vaccine-elicited mAbs. **(A)** Overview of the immunization and sampling of the rhesus macaques. **(B)** Immunization strategies and groups. **(C)** Example of B-cell sorting with 16055 NFL probes to identify mAbs with potent tier 2 autologous neutralization. **(D)** Neutralization activity (IC_50_) of mAbs isolated from each group. **(E)**. Antibodies isolated from various immunization trials and germline lineages. *,† Clonal variants

We used single memory B cell sorting to isolate mAbs from animals that showed the highest serum neutralization: 3 mAbs from group A animal D11 (D11A.F2, D11A.B5, and D11A.F9) [25], 3 mAbs from group B animal D15 (D15.SF6 and D15.SD7) and animal D16 (VD16.2C10); and 6 mAbs from group C animal D19 (D19.PA8 and D19.PD8) and animal D20 (VD20.1C7, VD20.1F9, VD20.5A4 and GM9_TH8) using 16055 NFL trimer probes. GM9_TH8 was isolated from animal D20 at week 47 after 3 additional boosts (post 6) with WT 16055 NFL trimer [26]. Neutralization against 16055 pseudovirus was assessed with potencies ranging from 0.005-4 μg/ml (**Fig 1C and 1D**).

All D11A antibodies share the same heavy chain germline VH4_3T_S3452 and JH4*01 genes, as well as the same light chain germline VL6-2*01_S6633 and JL2*01 genes (**Fig 1E**). Antibodies isolated from animal D15, D15.SF6 and D15.SD7 share the same heavy and light chain germline genes: IGH1_2L, JH4*01, IGLV6-2*01 and JL2*01 (**Fig 1E**). D19-isolated antibodies, D19.PA8 and D19.PD8, use different heavy chains germline genes, VH3_2T_S2563, JH5-2*02 and VH3_4A_S7053, JH4*01, respectively. D19.PA8 shares the same light chain germline genes IGLV6-2*01 and JL2*01 as the D15-isolated mAbs described above, while D19.PD8 uses the light chain germline genes lib5lambda_1 and JL1*01. VD16.2C10 uses the VH4.34_S2253, JH4*02 or JH5-1*02, lib4kappa_12 and JK2*01 germline genes. D20-isolated mAbs are clonally related and use the VH4.11_S9546, JH 5-1*01_S8786, lib4kappa_12 and JK2*01 germline genes [26] (**Fig 1E**). The somatic hyper mutation (SHM) levels in VH and VL range from 4.2-16.7% and 3.1-11.1% at the residue (aa) level, respectively (**Fig 1E**) [27].

### Vaccine-elicited antibodies interact with the V2b hypervariable region

Cross-competition binding analysis between the NHP neutralizing mAbs and known bNabs targeting different Env regions indicated that they all generally mapped to the V2 apical region of the 16055 NFL trimer while also displaying complete self- and cross-inhibition, assigning them to the same competition group (**S1B Fig**). Binding to 16055 gp120 constructs containing mutations in the V1/V2 loops confirmed specificity to the V2 region (**S1C Fig**). Epitope specificity was further mapped by neutralization sensitivity against a panel of 16055 pseudovirus mutants with residues along the 16055 V2 mini-loop (i.e., ^182^VPLEEERKGN^187^) mutated to alanine, or N187 mutated to glutamine (**S1D Fig**). The focused alanine scan confirmed dependence to the V2 hypervariable region as point mutants between residues V182 and K186C abrogated neutralization activity, while removal of the N187 glycan enhanced potency of the NHP mAbs (**S1D Fig**).

### Structural basis for HIV-1 tier 2 autologous neutralization

To understand the molecular basis for the tier 2 autologous neutralization from the isolated mAbs, we used a combination of negative stain EM (nsEM) and X-ray crystallography. To increase our chances of obtaining structural information, we used variational crystallography [28] where antigen binding fragments (Fabs) from a select number of antibodies complexed with 16055 NFL trimer, a scaffolded 16055 V1V2-1FD6 [29] or a 16055 V2b peptide [25] were purified and used for crystallization.

#### Group A mAb structural characterization

The nsEM of 16055 NFL trimer in complex with D11A.F9 (group A) and 35022 Fab [30] confirmed that the D11A.F9 approached its epitope located at the apex of the HIV-1 trimer horizontally, or parallel to the viral membrane (**Fig 2A**), consistent with previous studies [25]. D11A.F9 Fab crystals were obtained in complex with 16055 NFL trimer and 35022scFv [19, 31, 32], which diffracted X-ray to 6.5 Å. The low-resolution structure fitted well in the nsEM 3D reconstruction, confirming the horizontal angle of approach and (**Fig 2A**).

**Fig 2.**
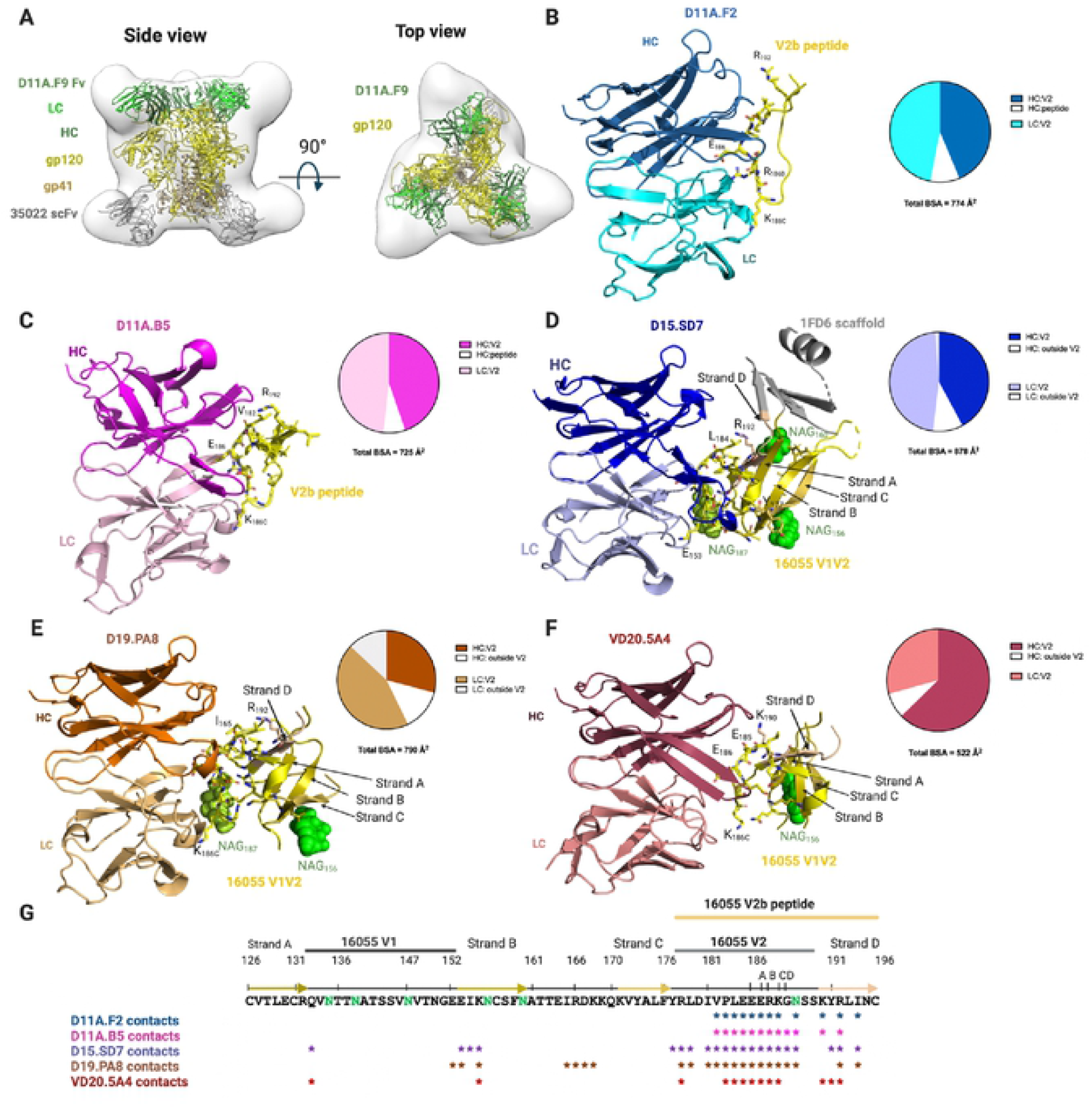
Vaccine-elicited antibodies recognize the V2 region of 16055 strain. **(A)** nsEM 3D reconstruction of the 1605 NFL/D11A.F9Fab/35O22Fab complex with low resolution crystal structure of D11A.F9 Fab (Heavy chain, dark green; Light chain, light green, Fv only is shown) and 35022 scFv (gray) in complex with 16055 NFL (gp120. yellow; gp41, light brown) shown in ribbon and in two different views. **(B, C)** Structures of D11A.F2 Fab (Heavy chain, sky blue; Light chain, cyan) and D11A.B5 Fab (Heavy chain, magenta; Light chain, light pink) bound to the V2b peptide (yellow). **(D)** Structures of D15.SD7 (Heavy chain, blue: Light chain, light blue). **(E)** D19.PAX (Heavy chain, orange; Light chain, light orange) and **(F)** VD20.5A4 (Heavy chain, raspberry; Light chain, light raspberry) Fabs in complex with tire 16055 V1V2-1FD6 scaffold. **(B, C, D, E, F)** Interacting residues are shown in sticks and glycans in green. Pie charts summarize the buried surface area (BSA) of the V2b and V1V2-1FD6. **(G)** Sequence of 16055 V1V2 highlighting the V2b peptide used for crystallization, the location of the V1, V2 and strands. Residues that contact the mAbs (within 5Å) are shown with asterisks underneath the sequence. N-linked glycosylation sites are shown in green.

Crystals of D11A.F2 and D11A.B5 were also obtained in complex with a 16055 V2 peptide, named here V2b peptide, ^178^RLDIVPLEEERKGNSSKYRLINC^196^ (numbering follows HXBc2 [33]), which diffracted X-rays to 2.8 Å and 2.0 Å resolution, respectively (**S1-3 Tables**). In both structures, the V2b peptide structure was fully resolved **(Fig 2B and 2C)** and adopted the same conformation as seen in the 16055 NFL trimer structure (RMSD of 1.1 Å and 0.8 Å over 17 and 16 Cα atoms, respectively) [18]. The high-resolution structures indicated that both D11A.F2 and D11A.B5 bind mainly to the 16055 V2 region, of which residues ^185^EEER^186a^ appear unique to the 16055 strain (**Fig 2B and 2C**). The D11A.F2 antibody buries ~ 774 Å^2^ of the V2b peptide, with ~701 Å^2^ in the V2 region and ~73 Å^2^ in Strand D [19, 29] (**Fig 2B, S2**). Similarly, D11A.B5 buries ~ 725 Å^2^ of the V2b peptide, with ~674 Å^2^ in the V2 region and ~51 Å^2^ in Strand D (**Fig 2C, S3**). We note that in both crystal structures, a shorter region of another V2b peptide appears to make additional interactions with D11A.F2 and D11A.B5. Since both the nsEM data and low-resolution crystal structure of D11A.F9 with 16055 NFL identified the hypervariable region V2 to be the epitope for D11A antibodies, we believe these additional contacts are not biologically relevant but the results of crystallization artifacts.

#### Group B and C mAb structural characterization

Crystals of D15.SD7 (group B), D19.PA8 and VD20.5A4 (group C) with the scaffolded 16055 V1V2-1FD6 were obtained and diffracted X-rays to resolution of 2.8 Å, 2.0 Å, and 2.8 Å, respectively (**Fig 2D-G, S1, S4-6 Tables**). The V1V2 structure adopts the same conformation as seen in the 16055 NFL trimer (RMSD of 0.9 Å, 0.9 Å, and 0.8 Å over 44, 43, and 41 Cα atoms, respectively), confirming that these antibodies recognize an epitope elicited by the trimer. Our structural analysis indicated that there are two copies in the asymmetric unit of D15.SD7/1FD6-V1V2 and D19.PA8/1FD6-V1V2 structures (**S4-5 Tables**). We observed clear density for three glycans in the gp120 V1V2 region at N156, N160 and N187 in one complex of D15.SD7/1FD6-V1V2, while the other complex in the asymmetric unit showed density for the N156 glycan only and thus chose the former for further analysis. Of note, D15.SD7 and D19.PA8 heavy chains showed some interactions with the 1FD6 scaffold (**S4-S5 Tables**), which we did not include in our analysis since they are not biological relevant. Additionally, the 1FD6 scaffold was mostly disordered in the D19.PA8 and VD20.5A4 complex structures (**Fig 2**).

Similar to the D11A antibodies, D15.SD7, D19.PA8, and VD20.5A4 bind mostly the V2 hypervariable region. They bury ~ 878 Å^2^, ~790 Å^2^ and ~522 Å^2^ of the V1V2, respectively (**Fig 2D-G**) of which ~781 Å^2^, ~579 Å^2^ and ~480 Å^2^ are in the V2 region only.

In conclusion, the structural analyses support our previous alanine scanning results, which showed that Glu^185^, Glu^186^, Glu^186A^, Arg^186B^, and Lys^186C^ mutations resulted in decrease or loss of neutralizing activities of D11A.F2 and GM9_TH8 [25, 26]. Indeed, all mAbs interact with the above-mentioned V2 residues (**Fig 2, S2-6 Tables**). We also observed additional interactions of all the mAbs with Val^182^, Pro^183^, Leu^184^, Gly^186D^, and Asn^187^ (**Fig 2, S2-6 Tables**), with the light chains of D15.SD7 and D19.PA8 showing some contacts with the proximal N-acetylglucosamine (NAG) at residue N187 (**Fig 2D and 2E**). We could not explain the slight difference in specificity at residues Pro^183^ and Leu^184^ described previously (Phad et al., 2020).

Finally, we also note that D15.SD7, D19.PA8 and VD20.5A4 all contact Lys^155^ in strand B as well as make additional contacts with the scaffolded V1V2 outside of V2 (**Fig 2**).

Since our high-resolution structures were solved with V2 peptide or V1V2 domain, we superimposed the above-described structures of mAb/V2b or V1V2 onto the structure of the 16055 NFL trimer (PDB ID: 5UM8) [19] by aligning the V2 or V1V2 region (**S2 Fig**). We observed that some mAbs showed additional contacts to the trimer not observed in our structures, either because the residues were not present in the V2 peptide and V1V2 domains or because these residues were disordered or did not show interactions in the solved structure (**S2 Fig**). Interestingly, in the superposition, mAb VD20.5A4 did not show additional contacts to the 16055 NFL trimer.

### Polyclonal antibody response to a similar epitope

Since mAbs elicited from vaccination target the same V2 region, unique to 16055, but used diverse germline genes and their autologous neutralization potencies differed by more than 1000-fold (IC_50_ ranging from 0.005 ug/mL (VD20.5A4, group C) to 3.68 ug/mL (D11A, group A)) (**Fig 1D**), we looked at differences and similarities of the paratope at the molecular level (**Fig 3**). We also analyzed the antibodies’ binding properties, including buried surface area (BSA), number of hydrogen bonds and salt bridges formed with the epitope, CDRH3 usage, electrostatics and angles of approach to decipher if some properties correlated with autologous neutralization potency (**Fig 4-7**).

**Fig 3.**
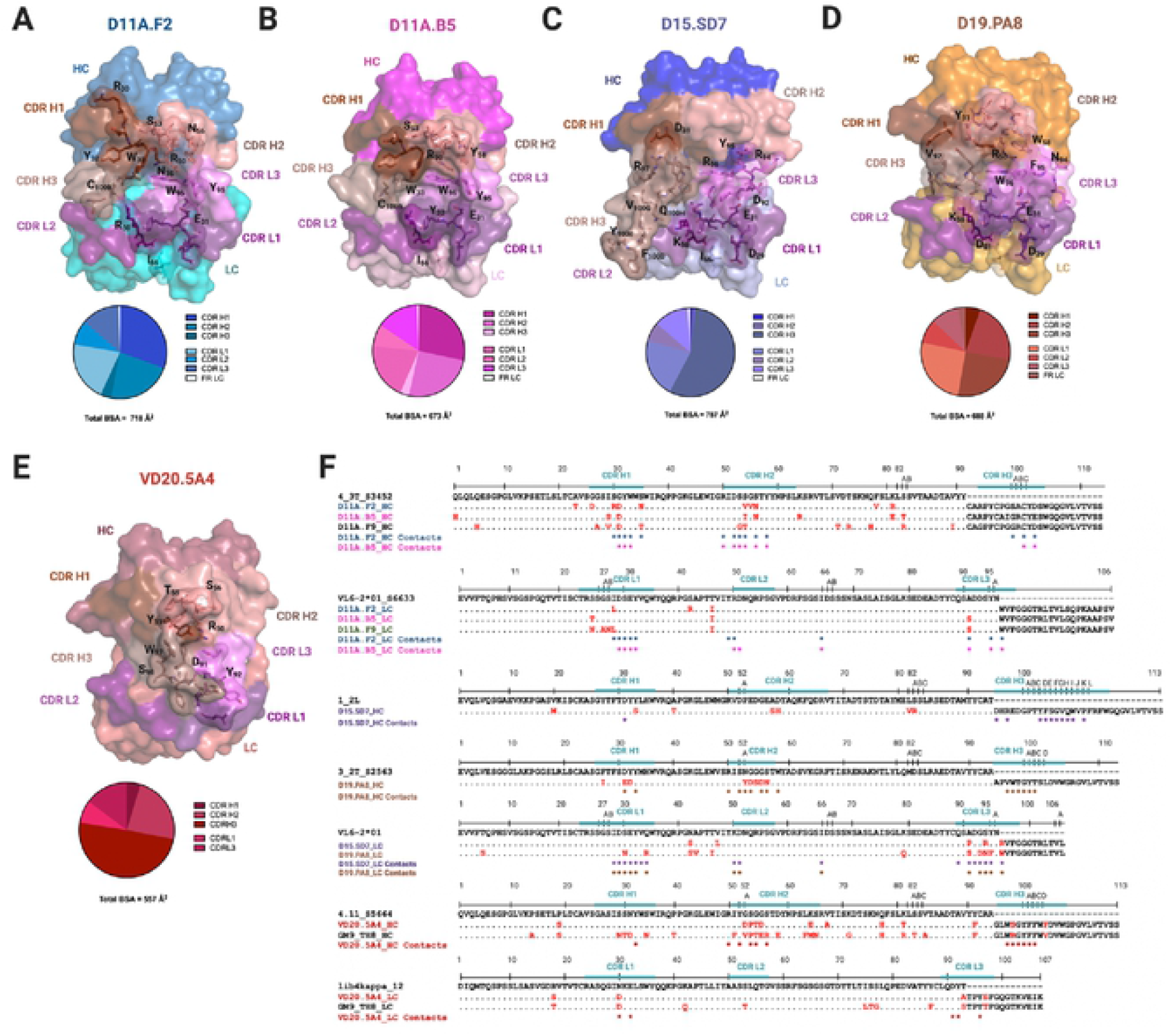
Structural characterization of mAbs elicited from vaccination. Surface representations of **(A)** D11A.F2, **(B)** D11A.B5, **(C)** D15.SD7, **(D)** D19.PA8, and **(E)** VD20.5A4 Fabs. All mAbs are color coded as follows: CDR H1, chocolate; CDR H2, salmon; CDR H3, dark salmon; CDR L1, violet purple; CDR L2, deep purple and CDR L3, violet. The pie chart represents the relative contribution of each CDR loops to the total buried surface area of the paratope for each mAbs. **(F)** Sequence alignment of the mAbs to their germline genes with CDRs highlighted. Somatic hyper mutations (SHMs) are highlighted in red. Residues interacting with 16055 V2b peptide or 16055 V1V2 are shown as asterisks below the sequence (contact residues within 5 Å).

**Fig 4.**
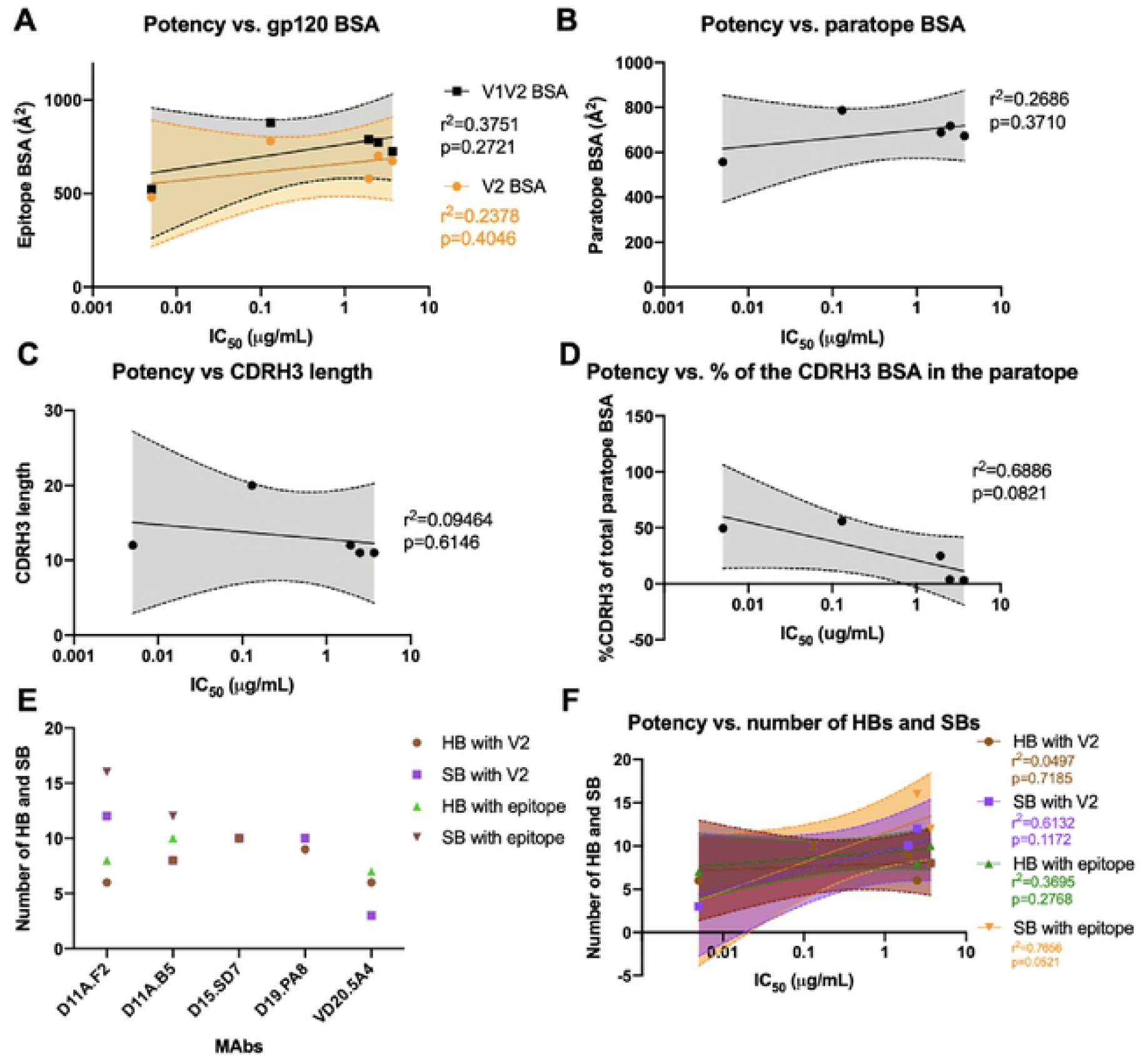
MAbs binding properties and correlations with autologous neutralization potency. Correlation between autologous neutralization potency and **(A)** epitope surface area, **(B)** paratope surface area. **(C)** CDRH3 length. **(D)** relative contribution of the CDRH3 surface area in the paratope and **(F)** number of Hydrogen Bonds (HBs) and Salt Bridges (SBs). The lines indicate the fitted linear regression model with 95% confidence shown in shaded grey or color as indicated. The r^2^ and p values are displayed. **(E)** Graph indicates number of HBs and SBs between the epitope/paratope with the different mAbs

**Fig 5.**
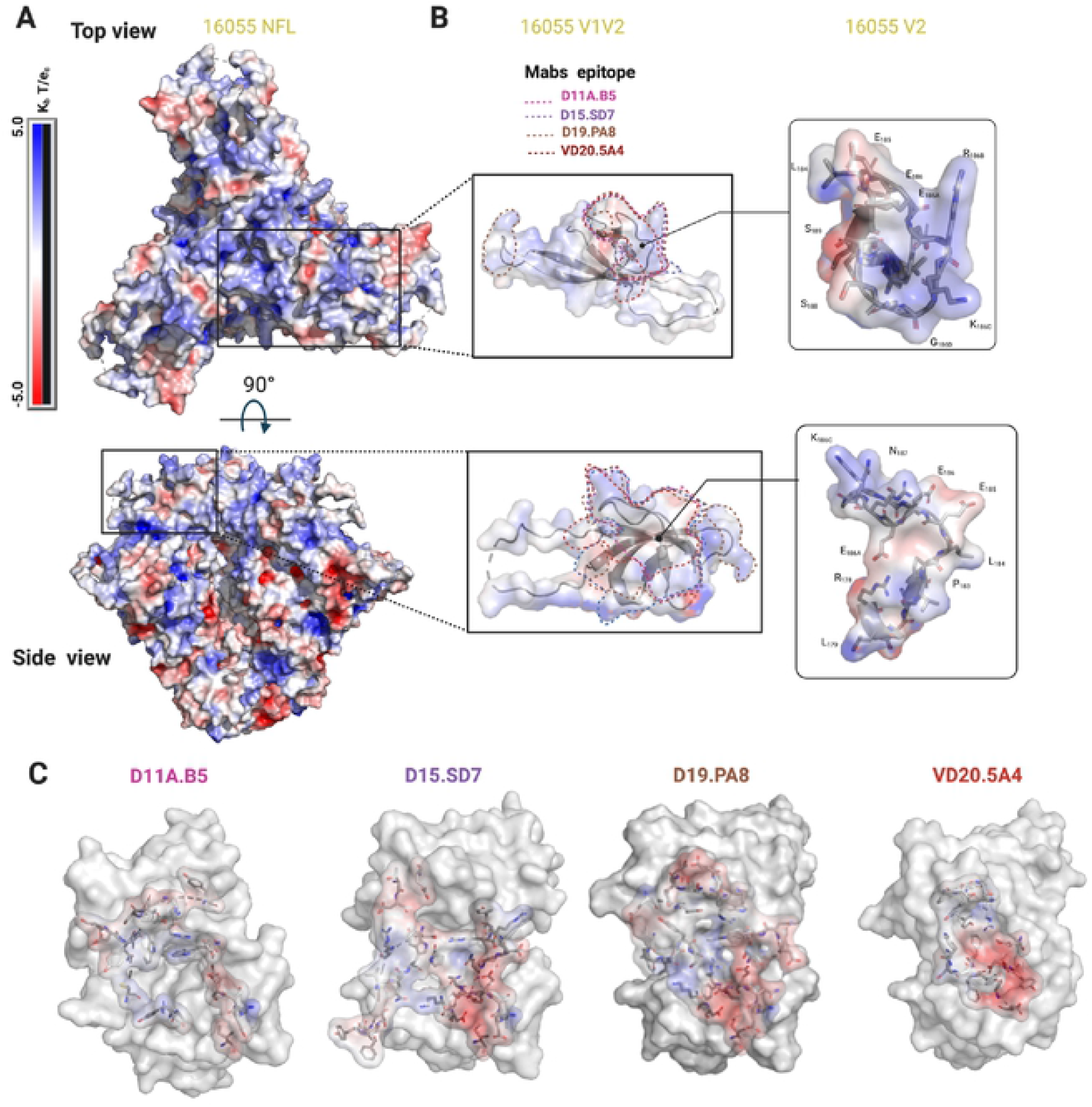
Electrostatics surface representation of 16055 NFL and different mAbs. Electrostatics surface representation of **(A)** 16055 NFL and **(B)** V1V2 and V2 region zooms (top and side views). Epitopes are highlighted by dotted lines and some residues in V2 are shown in stick and labeled. **(C)** Electrostatics surface representation of the mAbs paratope. Residues forming the paratope are shown in sticks.

**Fig 6.**
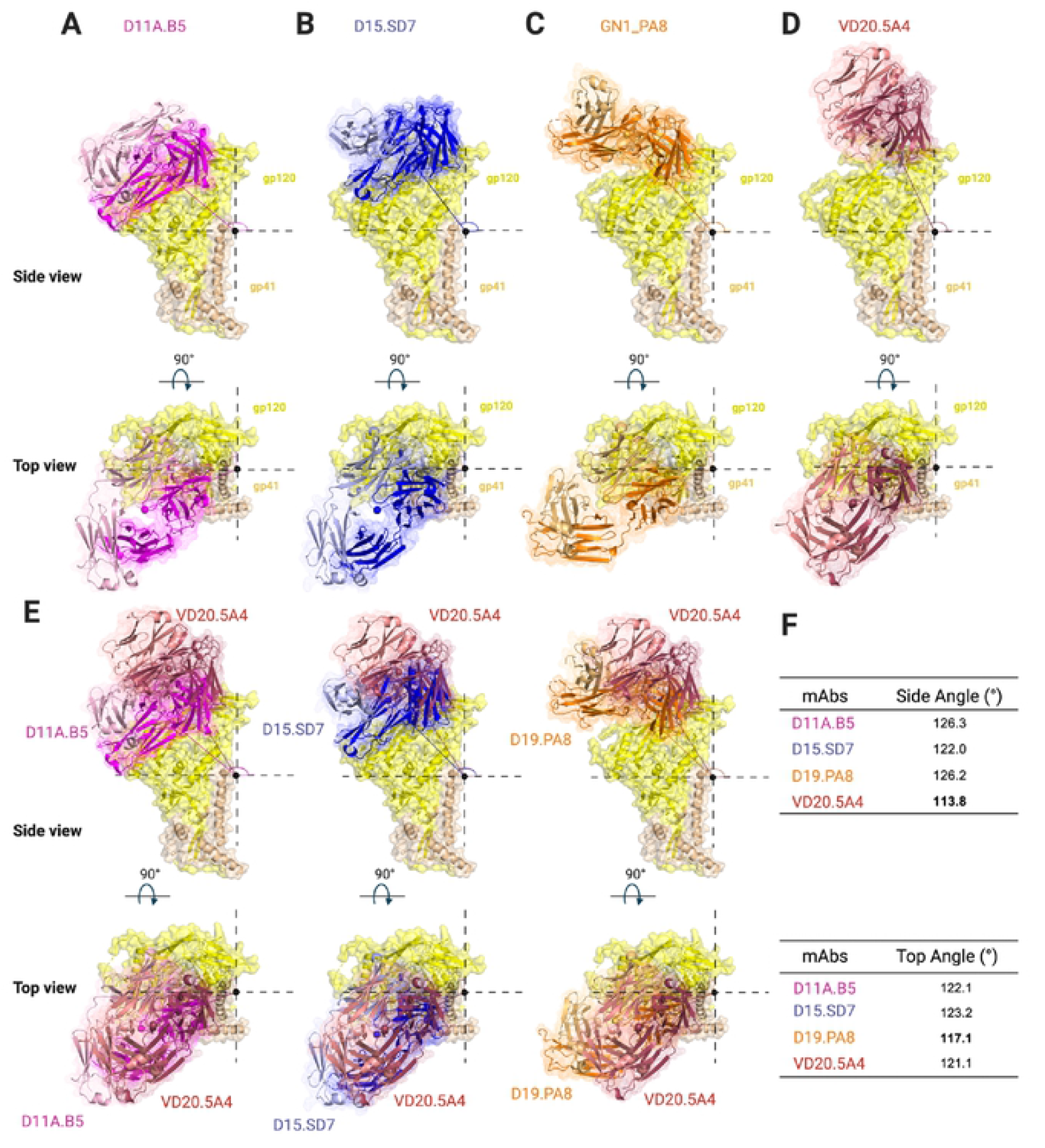
Vaccine-elicited mAbs target V2 region using a lateral approach with slightly different angles of approach. Side and top surface/cartoon representation of one gp120 (yellow)-gp41 (tan) 16055 NFL protomer with Fab bound: **(A)** D11A.B5 (pink), **(B)** D15.SD7 (blue), **(C)** D19.PA8 (orange) and **(D)** VD20.5A4 (raspberry) showing the angles of approach of each mAb. **(E)** Superimposition of gp120-gp41 16055 NFL protomer with D11A.B5 (pink), D15.SD7 (blue) and D19.PA8 (orange) onto VD20.5A4-bound gp120-gp41 protomer. **(F)** Summary of the mAbs angles of approach.

**Fig 7.**
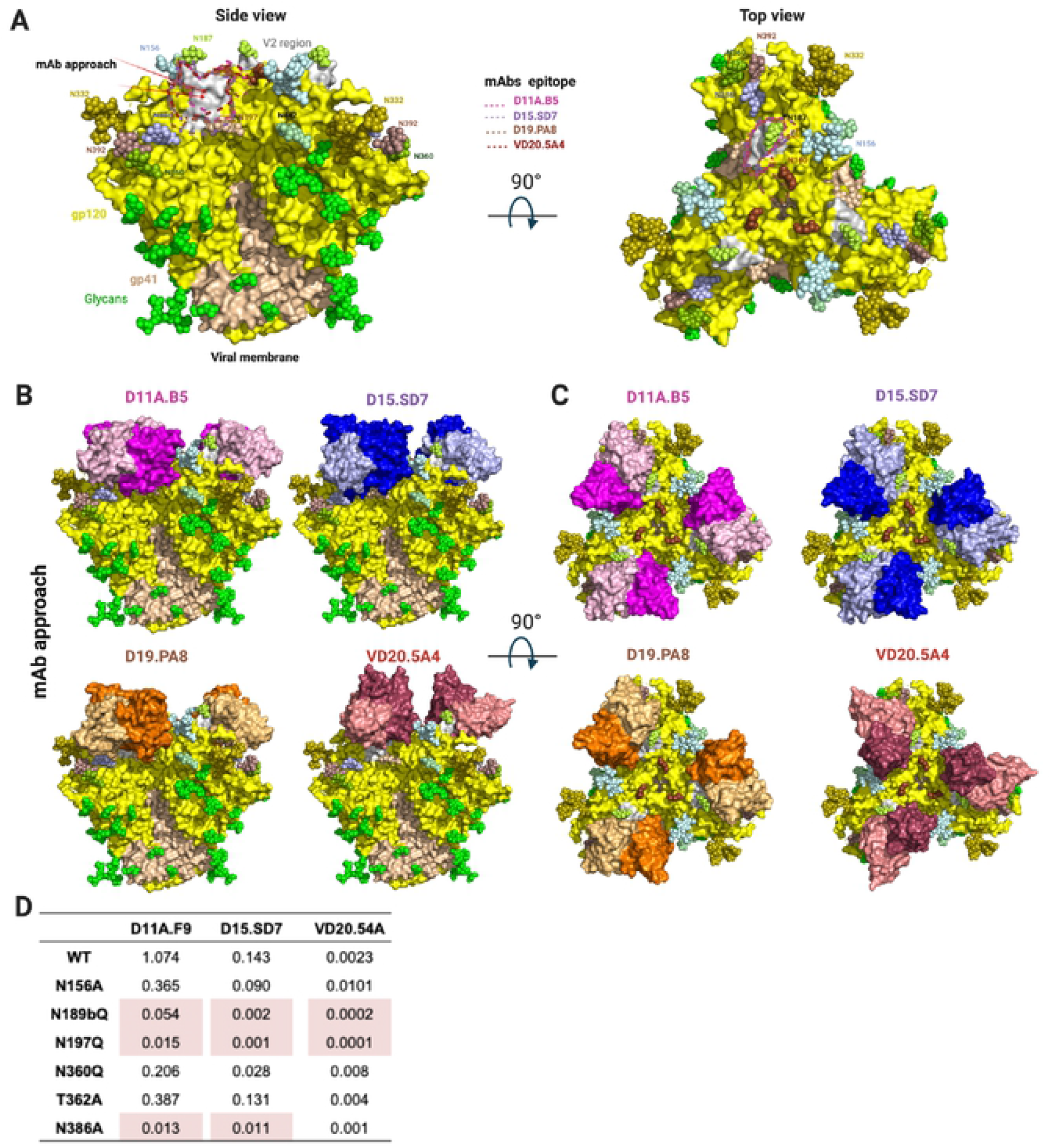
NHP Autologous Tier 2 neutralizing antibodies target a hole in the HIV-1 glycan shield. **(A)** Side and top view surface representation of 16055 NFL (PDB:5UM8) with gp120 shown in yellow, V2 region in grey, gp41 in wheat and glycans shown in green spheres or color-coded and labeled. Arrows indicate mAbs’ angle of approach. Epitopes targeted by the NHP mAbs are highlighted. **(B)** Side view and **(C)** Top view superpositions of the structures of D11A.B5, D15.SD7, D19.PA8 and VD20.5A4 onto the 16055 NFL trimer, showing how they access the glycan hole. Trimer and mAbs are shown in surface representation. Trimer is color coded as in **(A)** and mAbs as in Fig. 3. **(D)** Effect of glycan removal surrounding the epitope on neutralization potency. Neutralization IC_50_ values (μg/ml) shown with > 10-fold differences highlighted in red.

The total BSA of D11A.F2 is ~718 Å^2^, that of D11A.B5 is ~ 673 Å^2^, that of D15.SD7 is ~ 787 Å^2^, that of D19.PA8 is ~ 688 Å^2^ and ~ 557 Å^2^ of VD20.5A4 surface area is buried upon binding to its epitope (**Fig 3A-E**). We did not observe a correlation between the BSA of the paratope or that of the epitope with neutralization potency (**Fig 4A and 4B**). Indeed, VD20.5A4 is the most potent mAb but showed the least amount of BSA upon binding its epitope, indicating that in this case, precise targeting with a smaller epitope footprint might be relevant to potency.

The mAbs use all six complementary determining regions (CDRs) to bind their epitope, except for VD20.5A4, which does not use the CDRL2. D11A and GN1 mAbs also use part of the framework regions although these account for less than 6 % of the total BSA (**Fig 3**). Finally, both heavy and light chains are similarly involved in the interactions, except for D15.SD7 and VD20.5A4 which use primarily the heavy chain (58% and 78% of the total BSA paratope, respectively), with the CDRH3 accounting for 56% and 50% of the total BSA paratope and 97% and 64% of the heavy chain BSA, respectively (**Fig 3C, E, F**). The CDRH3 length varies from 11 to 20 residues but no correlation with potency was observed although D15.SD7 used primarily its 20-amino-acid CDRH3 to interact with its epitope (**Fig 4C**). We then assessed the correlation between potency and the relative contribution of the CDRH3 over the paratope (BSA from the CDRH3 over the total paratope BSA), and determined that there was a trend to significance correlation (**Fig 4D**). Indeed, it is interesting that the two mAbs that used most of their CDRH3 (in the context of our analysis, which only takes into account the V1V2 region and not the whole 16055 NFL trimer) proved to be the most potent autologous neutralizing mAbs.

All the mAbs used both germline and affinity matured V-gene residues in the interactions with their epitope, and within clonally related family, some of the interacting residues differ, however it is unclear what the difference or role in the affinity maturation is regarding the overall potency (**Fig 3F**). We note that the D11A mAbs have an intradisulfide bond in the CDRH3, which appears to rigidify the loop causing it to be less involved in the interactions. Such disulfide bonds have been observed before in mAbs isolated in humans with HIV and HCV infections [34, 35]. In these studies, the disulfide bonds were thought to be responsible for the antibodies’ neutralization potencies by stabilizing the affinity matured antibodies.

We next assessed the number of hydrogen bonds (HBs) and salt bridges (SBs) formed in each paratope/epitope interaction (**Fig 4E**). D11A.F2 and D11A.B5 form 8 and 10 HBs with the V2b peptide, 6 and 8 of which interact directly with the V2 region, respectively. In addition, D11A.F2 and D11A.B5 form 16 and 12 SBs with the V2b peptide, 12 and 8 of which interact with the V2 region, respectively. D15.SD7 and D19.PA8 form 10 HBs with their epitope, 10 and 9 of which interact with the V2 region, respectively. Moreover, D15.SD7 and D19.PA8 form 10 SBs with their epitope, all of them with the V2 region. VD20.5A4 forms 7 HBs (6 with the V2) and 3 SBs with the V2 region (**Fig 4E**). In conclusion, the number of HBs and SBs between the paratope/epitope did not correlate with the mAbs autologous neutralization potency (**Fig 4F**).

To further understand the differences in the potency, we looked at the electrostatics of the epitope and paratopes (**Fig 5**). While the epitope is overall positively charged (**Fig 5A and 5B**), the paratopes showed different electrostatics [36], with VD20.5A4 being strongly negatively charged towards the center of its paratope (**Fig 5C**). It appears that the paratope electrostatic of VD20.5A4 is more compatible with the overall positively charged epitope, which could explain its increased potency.

Finally, to understand the various mAbs’ angles of approach to their epitope on the 16055 NFL trimer, we superimposed the bound structures of D11A.B5, D15.SD7, D19.PA8, and VD20.5A4 on 16055 NFL trimer by aligning the V2b region of each structure to the NFL trimer (PDB:5UM8) [19] and calculated their angles of approach from a side and top view (**Fig 6**). Our analysis suggests that the D11A and GN1 mAbs approaches the V2 region with a similar angle (122-126°) from the side (lateral) (**Fig 6A-C, E-F**) while VD20.5A4 approaches the 16055 NFL trimer slightly from above (~114°) and rotated compared to the other mAbs (**Fig 6D-F**). While all mAbs approach their epitope with the same angle as seen from a top view, D19.PA8 is tilted 5° from the others (**Fig 6C and 6E-F**).

In conclusion, our structural analysis suggests that the difference in potency between the vaccine-elicited mAbs that bind the same epitope is likely due to differences in electrostatics in the paratope and angle of approach of the mAbs. Additionally, the nature of the CDRH3 interaction with the epitope also plays a role in the mAbs potency.

### Autologous tier 2 antibodies penetrate through the glycan shield

N-linked glycans extensively shield the surface of the HIV Env [13, 37] and this glycan shield is one of the reasons for Env’s resistance to mAb-directed neutralization [38–40]. Here, to explore the role of the glycan shield, we superimposed the structures of the vaccine-elicited mAbs in complex with their epitope onto the high-resolution structure of 16055 NFL trimer with N-linked glycans [19] (**Fig 7**). Interestingly, it appears that the mAbs target a partial glycan hole at the apex of the trimer formed by the long variable V2 sequence surrounded by glycans at position N156, N187, N197 and N386 (**Fig 7A**). The superposition also indicates that group A (D11A) and group B (GN1) mAbs will likely interact with glycans N197 (this one appears to clash in the superposition indicating that it must move out of the way for these mAbs to bind) and N386, while no glycans are in the way of group C antibody VD20.5A4. The conserved N-linked glycosylation site at residue N187 in the variable V2 region makes minimal contacts with the light chains of D15.SD7 and D19.PA8 (**Fig 2 and S4-5 Tables**), since there is electron density for the first proximal NAG in some molecules of the asymmetric unit. We performed site directed mutagenesis to evaluate the effect of removing certain glycans on neutralization potency.

In agreement with our structural observations, removal of glycan at position N386 enhanced the potency of D11A and GN1 mAbs with no effect to VD20.5A4. Additionally, removal of glycans at N187 and N197 showed increased potency (>10 fold) for all the mAbs (**Fig 7D**).

## Discussion

Revealing the molecular mechanisms by which vaccine-elicited antibodies target and neutralize the HIV-1 Env are invaluable to guide immunogen design and vaccine development [41, 42]. Our analysis shows that immunizations in macaques of a well-ordered 16055-based NFL trimer immunogen elicited mAbs that neutralize the autologous virus by targeting a gap in the dense glycan shield from which a small hypervariable V2 loop is antibody accessible. Germline gene analysis revealed that clonally distinct antibodies can target this same region with up to 1000-fold difference in potency. Interestingly, the most potent antibody, VD20.5A4 was not the most somatically hypermutated antibody. Our structural analysis indicates that the contribution of the CDRH3, electrostatics complementarity and angle of approach are likely responsible for the difference in potency between these mAbs. Of interest, VD20.5A4, which showed a smaller epitope footprint by targeting the epitope slightly more vertically than laterally compared to the others, also avoided most of the surrounding N-linked glycans. We can use the information collected here for immunogen “redesign” for more optimal targeting of relevant neutralization determinants. Although this loop is hypervariable in both length and sequence in each strain, it is likely often Ab-targeted [43], generating escape and hyper variability in humans [44]. This may be similar in terms of a shield breach in BG505, resulting in a vaccine-elicited immunodominant autologous neutralization response to the BG505 in rabbits, directed to a relatively large “glycan hole” at N241/N289 present in those trimers [45, 46].

Other V2-directed antibodies have been reported and characterized by some as four epitope families (V2p, V2i, V2q, and V2qt) based on their epitope [47, 48]. Based on our analysis, it is unclear if the mAbs described here fit in any of these previously described families, although they appear to overlap with the V2i mAbs, which recognized a discountinous epitopes in V2 overlaping the α4β7 integrin binding site [49]. It is thus possible that the mAbs described here can effectively elicit Fc-mediated functions.

Here we show that multiple clonally distinct antibodies elicited by 16055-based NFL trimers targeted a unique immunodominant V2 sequence. Interestingly, all antibodies recognize the conformation of this site as observed in the trimer context, even when co-crystallized with peptides or scaffolded V1V2, which can sometimes adopt other conformations [50], indicating that the use of well-ordered native like trimer as immunogens is likely needed to elicit mAbs that will bind such conformation. The data suggest that to induce breadth, structure-guided modification of strain-restricted but immunodominant gaps in glycan shielding, either holes or protruding loops, may shift responses toward more cross-conserved sites to better elicit cross-neutralizing responses to less immunogenic recessed determinants.

## Contact for Reagent and Resource Sharing

Further information and requests for reagents should be directed to and will be fulfilled by the Lead Contact, Marie Pancera

## Materials and methods

### Ethics statement

The animal work was conducted with the approval of the regional Ethical Committee on Animal Experiments (Stockholms Norra Djurförsöksetiska Nämnd). All animal procedures were performed according to approved guidelines.

### Animals

Female rhesus macaques (*Macaca mulatta*) of Chinese origin, 4-10 years old, were housed at the Astrid Fagraeus Laboratory at Karolinska Institutet. Housing and care procedures complied with the provisions and general guidelines of the Swedish Board of Agriculture. The facility has been assigned an Animal Welfare Assurance number by the Office of Laboratory Animal Welfare (OLAW) at the National Institutes of Health (NIH). The macaques were housed in pairs in 4 m^3^ cages, enriched to give them possibility to express their physiological and behavioral needs. They were habituated to the housing conditions for more than 6 weeks before the start of the experiment and subjected to positive reinforcement training in order to reduce the stress associated with experimental procedures. All immunizations and blood samplings were performed under sedation with ketamine 10-15 mg/kg intramuscularly (i.m.) (Ketaminol 100 mg/ml, Intervet, Sweden). The macaques were weighed at each sampling. All animals were confirmed negative for simian immunodeficiency virus (SIV), simian T cell lymphotropic virus, simian retrovirus type D and simian Herpes B virus.

### Immunization and sampling

Rhesus macaques were divided into groups and inculcated with variants of NFL HIV-1 Env trimers derived from the tier 2 clade C 16055 strain [19, 25]. Group A was inoculated with liposome-conjugated NFL trimers, Group B with NFL trimers lacking four N-glycosylation sites at residues 276, 301,360, 463 (del4) and Group C was inoculated twice with the del4 NFL trimers and then boosted with NFL trimers containing all glycans. All vaccines were administered with Matrix-M adjuvant, which was added to the immunogen prior to inoculation. Blood samples were collected two weeks after each vaccine inoculation. MAbs were isolated from the different groups as follows: D11A.B5, D11A.F2, and D11A.F9 (Group A), D15.SF6, D15.SD7 and VD16.2C10 (Group B), D19.PA8, D19.PD8, VD20.1C7, VD20.1F9, VD20.5A4 and GM9_TH8 (Group C).

### Neutralization assays

Neutralization assays were performed using a single round infectious HIV-1 Env pseudovirus assay with TZM-bl target cells [51]. To determine the mAb concentration or plasma dilution that resulted in a 50% reduction in relative luciferace units (RLU), serial dilutions of the mAbs and the plasma were performed and the neutralization dose-response curves were fit by non-linear regression using a 5-parameter hill slope equation using the R statistical software package. Site-directed mutagenesis to generate Env mutants were performed via QuikChange (Agilent Technologies) per the manufacturer’s protocol.

### mAb binding analysis by ELISA

NHP mAbs were tested for binding against 16055 gp120 or gp120 V region deletion mutants as previously described [25]. The gp120 deletion mutants include: ΔV1V2 (126-197), ΔV1 (134-153), and ΔV2 (159-193) with residues replaced with GAG or GGSGG for ΔV2. The mAbs were tested for binding using MaxiSorp 96-well plates (Nalgene Nunc International) coated at 2 μg/ml with wt gp120 or gp120 V region deletion mutants in PBS at 4°C overnight. After incubation with blocking buffer (5% non-fat milk/PBS/0.1% Tween-20), the mAbs were added and incubated for 1 hour at 37°C. Binding was detected by secondary HRP-conjugated anti-human Fcγ (Jackson ImmunoResearch) at 1:10,000 for 1 hour. The signal was developed by addition of TMB substrate (Invitrogen) for 5 min, reactions were terminated with 1 N sulfuric acid, and the OD was read at 450 nm. Between each incubation step, the plates were washed six times with PBS containing 0.1% Tween. For cross-competition ELISAs, NHP mAbs were biotinylated using EZ-Link NHS-Biotin (Pierce Biotechnology, Thermo Scientific) per the manufacturer’s protocol. 16055 NFL trimers were captured on the ELISA plate by a mouse anti-His tag mAb (R&D Systems) coated at 2 μg/ml in PBS at 4°C overnight. Five-fold serial dilutions of various bNAbs and non-bNAbs were pre-incubated with the captured trimer at RT for 30 min prior to addition of the biotinylated mAbs at a concentration previously determined to give ~75% of the maximum binding signal (i.e. binding to trimer with no competitor present) for 60 min at RT. The bound biotinylated mAbs were detected using HRP-conjugated streptavidin (Sigma) and TMB substrate with the reaction stopped with 1 N sulfuric acid. Competition is expressed as percent inhibition where 0% was the absorbance measured with no inhibitor present.

### Protein expression and purification

#### Antibodies

All antibodies were expressed in HEK293E cell lines as the expression platform. Cells were grown to a density of 1 million cells/mL and transfected using 293 transfection free reagent (Millipore) mixed with equal ratios of heavy and light chain encoding plasmids with 250 μg of DNA per one liter of culture. Expression was allowed to take place for 6 days, rocking at 37 °C. Cells were spun down at 4,000 rpm for twenty minutes and the resulting supernatant filtered. For Fab constructs containing a C-terminal His-tag on the heavy chain, supernatants were incubated with Nickel resin (TakaRa™) overnight at 4 °C, washed with several column volumes of 150 mM NaCl, 5 mM HEPES pH 7.5, and 20 mM Imidazole. The bound protein was eluted with 150 mM NaCl, 5 mM HEPES pH 7.5, and 300 mM Imidazole. IgG constructs were purified by affinity chromatography using GoldBio Protein A resin (GOLD BIO™). Following gravity flow, the resin was washed with multiple column volumes of PBS. The bound protein was eluted using IgG elution buffer (Thermo Scientific). Following affinity chromatography, the resulting eluate from Nickel or Protein A resin was further purified by size exclusion chromatography equilibrated in 150 mM NaCl and 5 mM HEPES pH 7.5.

#### 1FD6 scaffold

The 16055 V1V2 1FD6 scaffold was expressed in 293S GNTI^-/-^ cells (Howard Hughes™), following the same expression protocol as above. The supernatant was incubated with Nickel resin (TakaRa™) overnight at 4 °C, washed with several column volumes of 150 mM NaCl, 5 mM HEPES pH 7.5, and 20 mM Imidazole. The bound protein was eluted with 150 mM NaCl, 5 mM HEPES pH 7.5, and 300 mM Imidazole.

#### 16055 NFL

Expression of the 16055 NFL was the same at above. After pelleting the cells, the resulting supernatant was incubated with Galanthus nivalis agglutinin resin (Vector Labs™) overnight at 4 °C. The resin was washed with 20 mM Tris pH 7.4, 100 mM NaCl, and 1 mM EDTA, and bound protein was eluted with 20 mM Tris, 100 mM NaCl, 1 mM EDTA, 1mM methylmannopyranoside (MMP), pH 7.4 followed by further purification using SEC.

### Crystallization and X-ray data collection

#### Structures in complex with the v2b peptide

D11A.B5 and D11A.F2 constructs were mixed with the v2b peptide in a 2:1 peptide to antibody molar ratio and allowed to bind for one hour at room temperature. The complexes were screened against the Hampton Crystal HT, ProPlex HT-96, and Wizard Precipitant Synergy #2 crystallization screens. The NT8 robotic system (FORMULATRIX®) was used to set initial sitting drop crystallization trials. Following initial hits, crystallization conditions were optimized using hanging drop vapor diffusion. D11A.B5-v2b crystals were grown in 0.1 M Tris pH 8.0 and 1.32 M K/Na Tartrate at 8.5 mg/ml. Crystals were flash frozen in 120% of the crystallization solution supplemented with 15% 2R3R Butanediol. D11A.F2-v2b crystals were grown in 1.7 M Ammonium Sulfate and flash frozen in 2 M Ammonium Sulfate supplemented with 10% 2R3R Butanediol.

#### D11A.F9 and 35022scFv in complex with the 16055 NFL CC

The D11A.F9 and 35022scFv antibodies were incubated with the 16055 NFL at a three-fold molar excess of antibody. Complex formation occurred for one hour at room temperature. The complex was then treated for 30 minutes at 37 °C with the EndoH enzyme. Excess D11A.F9 and 35022scFv were purified away using an SEC column equilibrated in 150 mM NaCl and 5 mM HEPES pH 7.5. Final crystals were grown in 11% PEG 3350, 11% 2-propanol, and 0.1 M Tris pH 8.5, and flash frozen in 120% of the crystallization condition with 15% 2R3R Butanediol.

#### D15.SD7, D19.PA8 and VD20.5A4 in complex with IgG-16055 1FD6 scaffold

Typically, about 1.5 mgs of IgG was incubated with 1.5 mgs of 1FD6 scaffold for two hours at room temp before binding to Protein A resin. Unbound scaffold was removed by washing with 150 mM NaCl and 5 mM HEPES pH 7.5. HRV3C enzyme was added to generate Fab: scaffold complexes overnight at 4 °C. Following cleavage, the resulting flow thought was treated with EndoH for 30 minutes at 37 °C and then ran over SEC column equilibrated in 150mM NaCl and 5mM HEPES pH 7.5. Crystals were grown for data collection in the following conditions: The D15.SD7/V1V2-1FD6 crystals were grown in 0.2M Ammonium Sulfate, 0.1M MES pH 6.5, and 22% PEG 8000; D19.PA8/V1V2-1FD6 crystals were grown in 0.1M Tris pH 8.5 and 18% PEG 6000; VD20.5A4-1FD6 crystals were grown in 0.1M Na Acet pH 5.5, 10% w/v PEG 8000, 10% w/v PEG 1000, and 0.2M KSCN. Cryoprotectant solution was incorporated into the crystallization condition prior to crystal freezing.

#### Structure solution and model building

Data sets were processed using HKL2000, and initial models were generated using molecular replacement in Phenix. The D11A.F2-V2b, D11A.B5-V2b, D15.SD7-V1V2-1FD6 scaffold and D11A.F9 in complex with 16055 NFL and 35022scFv structures all used PDB 4RFO to search for initial molecular replacement solutions. PDB 5UTZ heavy chain and 4CQI light chain were used as molecular replacement search models for the D19.PA8/V1V2-1FD6 complex, and PDB 6U3Z was used as search models for the VD20.5A4-V1V2-1FD6 complex to find molecular replacement solutions. Following molecular replacement, iterative model building and refinement was achieved using COOT and Phenix, respectively. Summary of crystallization conditions for different complexes obtained.

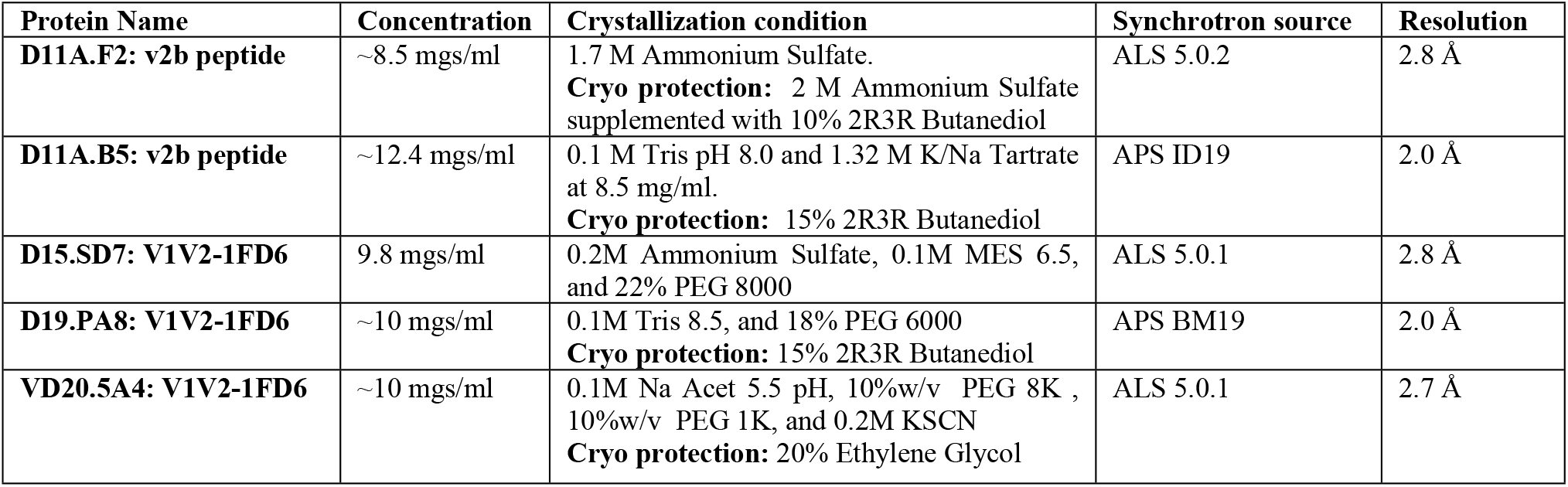

### Calculation of mAbs’ angles of approach

The V2 or V1V2 region from the structures of all antibodies complexes were first superposed onto the V1V2 region (residues 126 and 196 of gp120) of the 16055 NFL trimer (PDB ID: 5UM8). Chimera was used to determine the coordinates of the center of mass (COM) of the 16055 NFL trimer and each Fabs. The angles of approach of each mAbs were determined as follows: for the side angle approach, the X axis, the COM trimer and the COM of each Fab were used while for the angle of approach from the top the Z axis, the COM trimer and the COM of each Fab were used.

### nsEM

Complexes of 16055 NFL CC with three-fold molar excess of antibodies D11A.F9 and 35022scFv were prepared similar to that for crystallization studies. The samples were diluted to ~20 μg mL^−1^ and applied for 60 s to glow discharged Cu grids with continuous carbon film (300 mesh) (Electron Microscopy Sciences). Excess sample was blotted using a Whatman filter paper and stained for an additional 60 s using Nano-W (Nanoprobes). Excess liquid was blotted off and the grids air-dried for 1-2 minutes. Data were collected using an FEI Tecnai T12 transmission electron microscope operating at 120 keV. Images were taken using a Gatan 4Kx4K charge-coupled device (CCD) at a magnification of 67000X, corresponding to a pixel size of 1.6 Å, with exposure time of 1 s and defocus range of −1.0 to −2.0 μm. Single-particle EM reconstruction was performed using the Relion software package [52]. Particles were selected from 133 micrographs. CTF correction on the micrographs was carried out within the Relion software suite using CTFFIND [53]. A 4x binned stack of 47973 particles was created and subjected to reference-free 2D classification, and well-defined classes were selected. Selected particle images were then extracted as 2x binned set and subjected to 3D-refinement using a ligand-free structure of HIV Env (BG505.SOSIP) as initial model (PDB ID: 5ACO) [54]. This was followed by 3D classification of the particle images using the final structure from the 3D refinement as initial model. The best classes from 3D classification were grouped together to give a final set of 7685 particles. 3D refinement was performed again on this subset to give a final structure with a resolution of 15.75 Å.

## Data and Software Availability

The GenBank accession numbers for the novel neutralizing antibodies heavy and light chain of XXX-XXX reported in this paper are XXX-XXX. Coordinates and Structures factors have been deposited in the Protein Data Bank (PDB) under the PDB ID 6XLZ, 6VJN, 6WIT, 6WAS, and 6XSN for the antibody complexes. The crystals diffracted to high resolution at Structural Biology beamlines 5.0.1 and 5.0.2 and Argonne National Laboratory (ANL), Structural Biology Center (SBC) at the Advanced Photon Source (APS). Data reduction and processing were done using HKL2000, scaling with SCALEPACK, and phasing with PHASER using molecular replacement [55]. Model building was completed used Coot [56] and Phenix was utilized for refinement [57]. All structures were validated using MolProbity [58]. Structure visualization was done with Chimera [59] and PyMOL (The PyMOL Molecular Graphics System, Version 2.0 Schrödinger, LLC.). Figures were created using BioRender (https://app.biorender.com), Prism (GraphPad Prism version 9.0.1 for Mac, GraphPad Software, San Diego, California USA, www.graphpad.com), Chimera [59], and PyMOL (The PyMOL Molecular Graphics System, Version 2.0 Schrödinger, LLC.).

## Acknowledgments

We thank L. Stamatatos for use of laboratory space and equipment, Jason Gorman and Peter D. Kwong for providing the 16055 V1V2-1FD6 scaffold construct, the J. B. Pendleton Charitable Trust for its generous support of Formulatrix robotic instruments and an OctetRED384, Fondation Dormeur, Vaduz for generous support of equipment required for mAb isolation and Novavax, AB, Uppsala, Sweden, for generously making the Matrix-M™ adjuvant available to do this study. This project is funded by NIH grant P01 Al104722 to M.P., G.K.H., and R.T.W., and a Distinguished Professor grant (#2017-00968) from the Swedish Research Council to G.K.H., a Wenner-Gren Foundations fellowship to G.P.; NIH grants (R01 AI145055 and CHAVD UM1AI144462), Bill and Melinda Gates Foundation CAVD grant (OPP1084519), the IAVI Neutralizing Antibody Center, and the intramural research program of the Vaccine Research Center, NIAID, NIH. R01 AI145055 to R.T.W.; NIH grant number R01 AI140868 to K.K.L.; R.K. was supported by NIH T32 AI007509, and A.M. was supported by NIH T32 GM007750. Structural results shown in this study were collected at Structural Biology beamlines 5.0.1 and 5.0.2, which are supported in part by the National Institute of General Medical Sciences, National Institutes of Health. The Advanced Light Source is supported by the Director, Office of Science, Office of Basic Energy Sciences, of the United States Department of Energy under contract number DE-AC02-05CH11231. Also, part of results shown in this report are derived from work performed at Argonne National Laboratory (ANL), Structural Biology Center (SBC) at the Advanced Photon Source (APS), under U.S. Department of Energy, Office of Biological and Environmental Research contract DE-AC02-06CH11357.

## Author Contributions

Conceptualization: S.A., T.J.L., K.T., G.P., V.D., P.P., P. M-M., G.B.KH., R.T.W., M.P.

Formal Analysis and Software: S.A., K.T., G.P., G.B.KH. R.T.W., M.P.

Funding Acquisition: K.K.L., G.B.KH., R.T.W, M.P.

Investigation: S.A., T.J.L., K.T., G.P., A.M., R.K., V.M.P., K.K.L., G.B.KH., R.T.W., M.P. Methodology: S.A., T.J.L., K.T., G.P., S.S., V.D., P.P., P. M-M., J.R., R.K., S.OD., M.P.

Supervision: K.K.L., J.R.M, G.B.KH., R.T.W., M.P.

Writing – Original Draft: S.A., M.P.

Writing – Review & Editing: S.A., T.J.L., K.T., G.B.KH., R.T.W., M.P.

